# Context-Aware Technology Mapping in Genetic Design Automation

**DOI:** 10.1101/2022.08.24.505086

**Authors:** Nicolai Engelmann, Tobias Schwarz, Erik Kubaczka, Christian Hochberger, Heinz Koeppl

**Author notes:** The authors contributed equally to this research (Nicolai Engelmann, Tobias Schwarz, Erik Kubaczka).

## Abstract

Genetic design automation (GDA) tools hold promise to speed-up circuit design in synthetic biology. Their wide-spread adoption is hampered by their limited predictive power, resulting in frequent deviations between in-silico and in-vivo performance of a genetic circuit. Context-effects, i.e., the change in overall circuit functioning, due to the intracellular environment of the host and due to cross-talk among circuits components are believed to be a major source for the aforementioned deviations. Incorporating these effects in computational models of GDA tools is challenging but is expected to boost their predictive power, and hence, their deployment. Using fine-grained thermodynamic models of promoter activity we show in this work, how to account for two major components of cellular context effects: (i) Crosstalk due to limited specificity of used regulators and (ii) titration of circuit regulators to off-target binding sites on the host genome. We show, how we can compensate the incurred increase in computational complexity through dedicated branch-and-bound techniques during the technology mapping process. Using the synthesis of several combinational logic circuits based on Cello’s device library as a case study, we analyze the effect of different intensities and distributions of crosstalk on circuit performance and on the usability of a given device library.

## 1 Introduction

Although genetic design automation (GDA) has made significant progress in recent years (*1–3*), its wide-spread adoption among synthetic biology researchers is still hampered by the limited number and modest size of well-characterized part libraries but also by the limited predictive power of those tools. That is, the circuit designs found by employed computer models often do not operate as predicted when realized within a cell. There are several reasons for this but to some extent they can be traced back to the fact that our mechanistic understanding of the complex molecular biology operating within a cell is still incomplete. First, this leads to part libraries possibly omitting key physical and biochemical parameters that are required to fully specify the behavior of a part. As a consequence, the derived part models are under-specified (*4*). Second, the interactions among parts of a genetic circuit and moreover between those parts and the host cellular machinery are even less characterized or understood and not accounted for in current GDA tools. All those potential interactions are traditionally subsumed under the term context effects (*5, 6*). These effects include but are not limited to (i) the dependence of a genetic part on the adjacent up and downstream sequences (*7*), (ii) the cross-talk among synthetic regulators due to their limited binding specificity (*8*), similarly (iii) the titration of regulators to non-cognate binding sites on the host genome (*9*), (iv) the retroactive or loading effect exerted on upstream circuits elements by subsequent downstream loads (*10, 11*) and (v) the cross-talk between the host and genetic circuits on the energetic level through, e.g., the competing use of the energy-intensive translation machinery (*12*).

Molecular means to counteract detrimental context effects are devised on a case-by-case basis (*13–15*) but their systematic inception requires models deployed in GDA tools to account for those effects. Moreover, certain titration effects turned out to be beneficial and could be leveraged with GDA to shape gene response characteristics if used in a controlled manner (*16*).

Incorporating context effects into circuit models involves extending them through first principles biophysical considerations, through acquisition of appropriate characterization data (circuit behavior under different environments) and – most effective – through a combination of both. Context effects stemming from the dynamic molecular environment of a host cell will give rise to measurable cell-to-cell variability of circuit behavior (*17*). Those context variations were shown to be dominant contributors to noise, surpassing intrinsic contributions due to small-copy-number fluctuations (*18*). Hence, single-cell data for parts or circuits can be used to learn or calibrate context models. In turn, GDA tools incorporating context-effects can then evaluate and score a circuit candidate based on its generated single-cell data across a in-silico cell population (*19, 20*). This provides means to identify circuits candidates or additional regulatory motifs leading to robust circuit behavior in the presence of context-effects.

Among the many context-effects, two of them can be tackled by relying on available, more detailed biophysical models of gene regulation. That is, (i) cross-talk among gene expression units stemming from limited TF binding specificity of deployed transcription factors and (ii) titration of those TFs to non-cognate binding site among the host genome can be dealt with through thermodynamic models developed over the last two decades (*21, 22*). They work out the statistical mechanics of promoter occupancy states in order to derive rates of gene expression in nearly parameter-free manner. They have been particularly successful in predicting promoter strengths within procaryotic cells, where this occupancy dynamics is believed to happen at thermodynamic equilibrium. Moreover, they allowed to quantitively reproduce the effect of titrating DNA binding sites on the dose-response characteristics of repressors in E. coli (*9*).

Cello (*2, 23*) is a well known tool for GDA. Nevertheless, its circuit scoring function does not take into account context effects. It rather uses the scoring function to heuristically optimize the assignment of library gates to the gates of a circuit. In a similar way iBioSim (*24*) is a GDA tool that employs branch and bound to optimize the gate assignment. The optimization goal in this tool is the length of base pairs. Thus, it does not ensure the proper realization of the wanted circuit function. Also, this tool does not take into account context effects. To the best of our knowledge, no current GDA tool is capable of optimizing a circuit realization under context effects.

In this paper we present an approach for incorporating the effect of the two aforementioned context-effects in the GDA technology mapping process. In particular, we make use of and extend thermodynamic models for gene regulation. As a consequence, the gates within the device library have a more fine-grained computational representation and ad-hoc phenomenological Hill-type characteristics can be avoided. As the incorporation of cross-talk renders the considered class of combinational circuits to be effectively sequential circuits, i.e., dynamic circuits encompassing feedback loops, we discuss the computational consequences of this and lay out efficient algorithms. In order to overcome the increased computational complexity incurred by such feedback structures, by the fine grained thermodynamic models, and by the single-cell circuit scoring we devise an efficient branch-and-bound algorithm for the mapping process. In particular, we derive score intervals for partial circuit topologies and based on them allow to prune entire branches of candidate assignments of library gates without evaluation even in the presence of cross-talk. We demonstrate the methodology using the gate library of Cello presented in (*23*), where we artificially introduced non-cognate binding affinities of TFs for parts within the circuit and for the genomic DNA of the host.

## 2 Results and Discussion

### 2.1 Logic Circuit Architecture

The circuit and gate architecture in this work is similar to the one used in the Cello suite (*23*). Therefore, combinational logic circuits are built using NOR logic. This implies that NOR gates are the only type of logic gate allowed in the circuit. The constituting set of parts comprising a single NOR gate consists mainly of a fixed combination of a gene and an output promoter. The gene expresses a repressing transcription factor (TF) which binds to its cognate binding site at the output promoter. The output promoter activity then acts as the output level of the gate. Promoter activities of (non-cognate) input promoters placed upstream (in the sense of a transcription direction) of the gene act as input levels of the gate. Thus, the TF expressed by the gene connects a gate’s input with its output in the form of a gate-internal connection. Binding characteristics of the TF with its cognate output promoter and expression characteristics of the gene then implement the gate’s transfer function.

### 2.2 Modelling Context-Effects in Genetic Circuits

#### 2.2.1 Nuisance-free Thermodynamic Circuit Description

Modelling the NOR gate’s transfer functions using equilibrium thermodynamics allows convenient integration of the considered context effects. We will elaborate on this in the following.

We start with a given concentration *f* of a translated product, i.e. its copy number w.r.t. constant volume. This product can be e.g. a TF or a reporter. Since we work with averages of statistical ensembles, we also introduce an associated random variable *F* for the product concentration. Another random variable *X* ∈ {0, 1} accounts for the occupation of a single promoter upstream of the product’s encoding gene by RNA-Polymerase (RNAP). Finally, a collection of random variables *C* is used to quantify any considered molecular context moderating RNAP occupation of the promoter, i.e. changing *X*’s distribution. Now, let *F* = *f* be the realization of the product concentration given a fixed molecular context *C* = *c*. A fundamental model assumption w.r.t. gene expression is then that f and the expectation E (*X* | *C* = *c*) are proportional, i.e. *f* ∝ E (*X* | *C* = *c*). This expectation E (*X* | *C* = *c*) is termed occupancy. A second fundamental model assumption directly related to equilibrium thermodynamics is that single binding states, i.e. the permutations of assignments of available molecules to available binding sites, follow a Boltzmann distribution. This itself implies that energetically indistinguishable binding states are equally probable. Thus, the model reduces the underlying complicated mechanics of gene expression to combinatorics on the ensemble of RNAP binding states (*21*). Naturally, quantities considered to moderate RNAP promoter occupation (e.g. TF’s) and thus included in C either inflate the ensemble or increase statistical dependence among the binding states and thus complicate the combinatorics (*25*). Approximate expressions then need to be found that are reduced to the dominating terms to maintain a certain scalability. As a final assumption, we set the promoter activity *α* to be proportional to the promoter occupancy, i.e. *α* ∝ E (*X* | *C* = *c*), and thus *α* ∝ *f*. An illustration summing up the interactions that we can tackle using the equilibrium thermodynamic model is shown in Fig. 1A.

**Figure 1:**
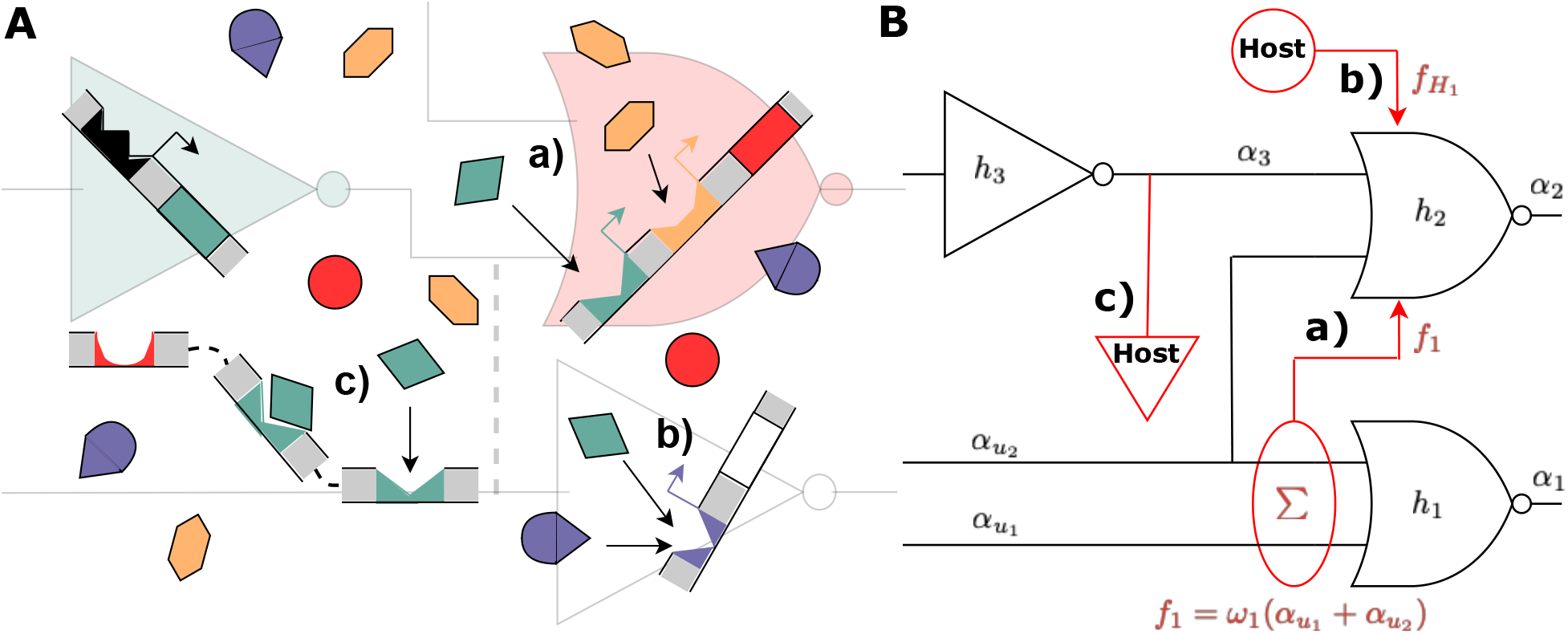
*A* Illustration of the thermodynamic perspective of the logic circuit. a) desired binding of cognate TFs to cognate promoters implementing the gate’s transfer function. b) crosstalk with non-cognate TF binding amplifying repression at the non-cognate promoter. c) titration of available TFs to the host genome or other non-specific binding sites; *B* Circuit topological view of the effects available to the model. a) crosstalk from another gate’s internal TF. b) crosstalk from out-of-circuit TFs available through the host context. c) titration to non-specific binding sites available through the host context.

To initiate construction of a genetic logic circuit using thermodynamic gene expression, we first consider the construction of a simple NOT gate. We can understand this NOT gate as a NOR gate with a single input. As described in section 2.1, the gate is comprised of an output promoter, its cognate (gate-internal) TF, and the gene that encodes the TF. To obtain the full transfer function *h*, we need to give it an input promoter activity *α*_in_. The transfer function h determines the output promoter activity *α* as a function of its input promoter activity *α*_in_. Let’s say, the cognate TF concentration f is known and there is no cooperativity. Then, the promoter activity *α* can be given in terms of *f* by

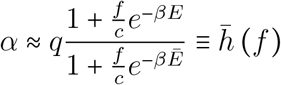

where *β* ≡ (k*T*)^-1^ is a thermodynamic constant with *k* the Boltzmann constant and *T* the temperature. *E* is a relative energy term encoding the energy expense of having an RNAP and a TF bound at the promoter. 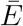 on the other hand denotes the expense of having only a TF and no RNAP bound. The number *c* is the concentration of non-specific “background” binding sites that are assigned average binding energy and *q* is a proportionality constant. With the exception of *q*, a similar expression can also be found in (*26*). In this work, we consider cooperativity by recruitment. This means any bound TF reduces the energy expense of an additional TF to be bound as well. We assume this effect to saturate at around *N* bound TFs and thus introduce *N* as a “cutoff” order of cooperativity. As a consequence, we allow the relative energies *E*(*n*) and 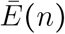 to obtain an integer argument encoding how many TF are bound. For promoter activity *α*, we then obtain the expression

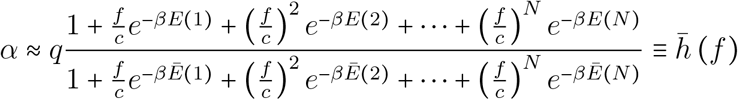

The last step is to formulate the *f* as a function of *α*_in_ to obtain the gate’s full transfer function *h*. Since *f* and *α*_in_ are proportional, we can express *f* ≈ *ωα*_in_ using the proportionality constant *ω* and finally obtain

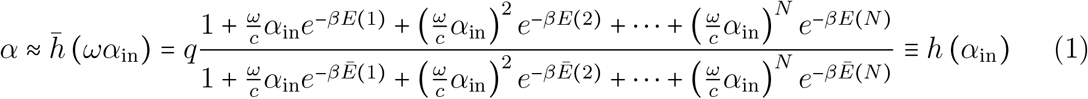

We show in Methods 4.1.2 that 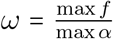. In the following, we introduce the circuit and in this context the expression for the transfer function of a NOR gate with an arbitrary number of inputs. Because of the complexity of the resulting expressions, we will resort to a more abstract notation in comparison to (1).

Let in the following the set of all gates in the circuit be 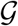 and we have a set of integers 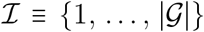 enumerating the gates. 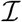 assigns an index to each gate, so that for any gate from 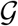 with index 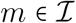, the quantity *f_m_* is the concentration of the gate’s affiliated (internal) TF expressed by the gate’s gene. Again, we assign a random variable *F_m_* to this concentration. The random variable of RNAP occupation of its output promoter will be written *X_m_* and the collection of random variables considered to comprise its relevant molecular context will be *C_m_*. The promoter activity of the output promoter of gate *m* is then denoted by *α_m_*. In this section, only the ideal case of no titration nor crosstalk is considered. Thus, besides gate independent host constants, we consider only the affiliated TF concentration *f_m_* to be relevant in moderating RNAP occupation of the output promoter, i.e. *C_m_* ≡{*F_m_*}.

Following the steps for the simple NOT gate, to obtain the NOR gate model, we first consider the function mapping the input promoter activity to the internal TF concentration *f_m_*. The input promoter activity is roughly given by the sum of activities of all promoters upstream. Let 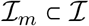 denote the set of indices of gates wired to the input of the gate with index *m*. Then,

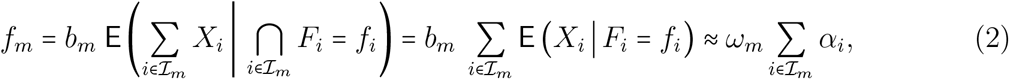

where *α_i_* is the promoter activity of the output promoter of the *i*-th gate posing as one of the *m*-th gate’s input promoters and *ω_m_* is a gate-specific proportionality constant. Second, we consider the promoter activity of the output promoter of the *m*-th gate as a function of the TF concentration *f_m_*. We again assume that there exists a “cutoff” integer *N_m_* which bounds the significant order of cooperativity for gate *m*’s TF. As a result, we obtain for the transfer function *h_m_* of the *m*-th gate, i.e. for its output promoter activity *α_m_*

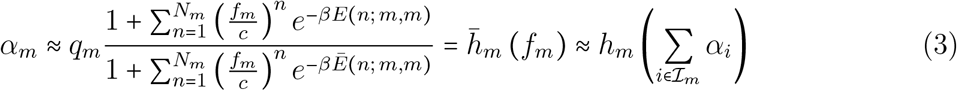

where *q_m_* is a proportionality constant. We again expanded the relative energy functions *E* and 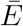 by two integer arguments *i* and *j*, so that *E*(*n*; *i,j*) is the relative energy of the binding state where *n* TFs from the *i*-th gate bind to the *j*-th gate’s binding site. Since we consider no crosstalk here, only the *E*(*n*; *i,i*) are occuring (because they are finite). Note, that also *α_m_* ∝ E (*X_m_*| *F_m_* = *f_m_*).

As a template model for the repression mechanics, we used a version of the “simple repression” from (*21*), a model suitable to describe most repression mechanisms in bacterial cells. This choice is reflected in the simple rational appearance of (3). To implement basal expression, we adopted the concept of imperfect competition between RNAPs and repressors from (*26*). This allows RNAPs to bind to promoters and initiate transcription under large energy expense even if a TF is bound to the repressor binding site. The original model prohibits this combination completely and is thus not able to model leaky gene expression in high copy number regimes of repressors. The choice to add this mechanism results in the appearance of the numerator polynomial featuring the larger energies E (in comparison to 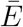) in (3).

As suggested by the two fundamental model assumptions and reflected in (3), the circuit can be fully parameterized using binding energies and proportionality constants - besides constants associated with the host environment that are independent of the circuit. Indeed, for the calculation of the whole circuit response, only these quantities are needed. In the following, we describe how to calculate this response.

Assume now, we are given a set of promoter activities 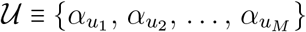 as circuit inputs for a whole circuit. We further choose a set 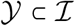 of gate indices referring to those gates whose output promoter activities pose as the circuit’s output. Since any *α_m_* for 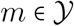 cannot be measured directly, we introduce a post-processing circuit that maps the output activities to a measurable reporter concentration, e.g. that of YFP. This post-processing circuit then implements the transfer function 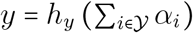 with the reporter concentration *y*. As an example, we refer to Cello (*23*) where *h_y_* is the identity function so that there 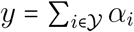. Then, the circuit response has a unique solution *y* for an input 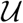 if and only if the equation system

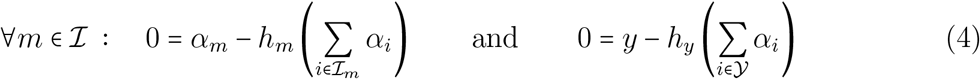

has a unique solution. In the case of a valid combinational circuit where each gate can be described by an equation of the type of (3), the system has a solution and there exists an order of successive calculation of (4) following which, each equation can be solved explicitly.

Using the thermodynamic model, we can also introduce titration effects with the host genome. In this work, only first-order titration effects are considered. Thus, given a set of off-target binding sites provided by the host environment that attract a specific TF, we consider titration of these TFs away from the circuit. We call these host-provided off-target binding sites competitor sites. Introduction of a concentration *s_m_* of competitor sites attracting the *m*-th TF can be effectively modelled using the transformation 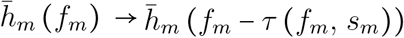 where *τ*(*f_m_, s_m_*) is a monotonically increasing nonlinear function of *f_m_* and *s_m_* that can be pre-computed for every fixed host-configuration. Then, we obtain for the modified transfer functions

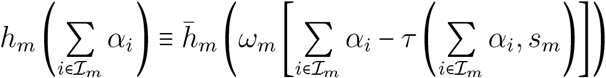

where the function mapping the TF concentrations to the output promoter activity 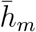 stays unchanged as in (3). Note, that this modification can be immediately applied to any gate for a fixed host and the circuit equation (4) stays formally unchanged using the modified *h_m_* and *h_y_*.

#### 2.2.2 Calibration to Cello’s Gate Library

To obtain meaningful parameters for a whole set of NOR gates, we adopted the gate library from the Cello suite (*23*) and calibrated our gate’s transfer functions to match the corresponding transfer functions from the Cello gate library as closely as possible. For this, we used Simulated Annealing to minimize the logarithmic quadratic error over 1000 collocation points equally spaced in the logarithmic domain within the interval [10^-4^,10^2^] w.r.t. the thermodynamic parameters. The obtained curves had a peak | average | least cumulative (integrated) error of ~ 0.193 | ~ 0.0166 | 10^-19^ in the logarithmic domain. To visually grasp this result, we supply the plots comparing both transfer functions in the supplementary material.

#### 2.2.3 Adding Crosstalk

The main advantage of the thermodynamic approach w.r.t. modelling crosstalk is the explicit formulation of TF binding. Therefore, to implement crosstalk for the target gate with index 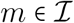 we need to consider the occupancy E (*X_m_* | *C_m_*) to be dependent on more than only the cognate TF {*X_m_*} ⊂ *C_m_*. In thiy general case, where we assume the binding specificity of any TF to be potentially imperfect, we need to consider the concentrations of all available TFs, so 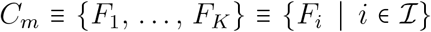. We then find for the output promoter activity 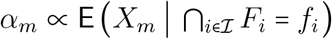 that

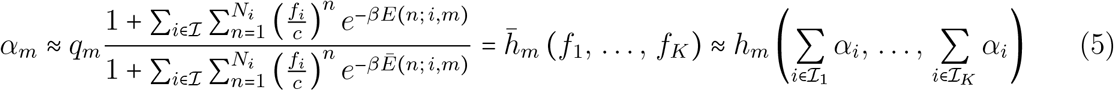

Again, similar to above the circuit response has a unique solution *y* for the input 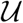 if and only if the equation system

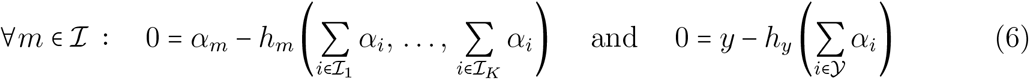

has a unique solution. In contrast to (4), equation (6) usually cannot be solved explicitly since the required order of successive calculation doesn’t exist. Since crosstalk in general introduces feedback connections to the circuit, the computational resources needed to solve (6) are comparable to those needed for asynchronous sequential circuits. Therefore, root finding or - in this case preferably - fixpoint algorithms need to be used to find a solution.

We explain how we solved (6) in Methods 4.2. In contrast to the combinational case (4), where each *h_m_*, 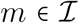 and *h_y_* need to be evaluated only once, we need to iterate over *h_m_* and *h_y_* a few times until convergence. While the amount of iterations greatly varies with the circuit topology, around 10-20 iterations have been observed in an average scenario. Thus, using our approach an increase in simulation runtime by one order of magnitude has to be expected under consideration of crosstalk. Note, that this increase depends on the algorithm used and we cannot rule out the possibility to reduce the amount of iterations significantly by choosing a specific solution algorithm.

This increased demand of computational resources to determine the circuit response under crosstalk aggravates the already present runtime bottleneck in technology mapping of genetic logic circuits. While technology mapping in EDA typically finds near-optimal assignments for huge circuit structures in amenable time, in GDA the pronounced quantitative differences of functionally similar parts - and thus, their limited mutual compatibility - make already finding better-than-average assignments in small circuit structures computationally expensive. To tackle this bottleneck, we present a novel technology mapping procedure specifically tailored to the optimization requirements in current GDA problems in the following section.

### 2.3 Efficient Context-Aware Design of Genetic Circuits

A main objective of the design of genetic logic circuits is finding a topology and an assignment of gates from a library to this topology that maximise a given performance metric. In contrast to EDA, where metrics typically are based on circuit speed and size, metrics in GDA take into a account whether a circuit fulfills its functional specification. For example, the GDA tool Cello uses a circuit score based on the separation of the complementary Boolean outputs (*23*). It has been shown that the circuit topology plays an important role in genetic circuit design (*19*). Yet, the computationally challenging part of circuit design is finding an assignment of genetic gates from a library to the topology. Each single assignment needs to be simulated in order to evaluate its score, making the gate assignment a combinatorial optimization problem. To this end, we propose a method based on the Branch and Bound (B&B) optimization approach that takes into account context effects. It leverages fundamental features of the dose-response curve of gates based on transcriptional regulation and finds the optimal solution with respect to Cello’s circuit score. First, we introduce a notion of signal-compatibility of genetic gates that integrates into the B&B scheme and ensures robustness with respect to variability.

#### 2.3.1 Compatibility of Genetic Gates

In contrast to electronics, genetic gates feature different transfer characteristics even if they represent the same logic function. This can lead to mismatches in the signal levels in a cascade of gates. In extreme cases, the complementary Boolean outputs of a logic gate can fall into the same region of a subsequent gate. This leads to a loss of logic information and thus to a non-functioning genetic circuit. In less severe cases, the subsequent gate is operating near the transition region of its transfer function. This results in a reduced tolerance with respect to signal perturbations and should be avoided when designing robust genetic circuits. In Cello, this is taken into account by ensuring that the minimum and maximum output values of a gate lead to an output of the subsequent gate that is higher than half the maximum or lower than twice the minimum output value (*23*). Circuits that violate this criterion are filtered out during the technology mapping process. We propose to generalize this concept by defining 3 dB-thresholds of the output signal of each gate and determining the corresponding input signal levels (see Fig. 2). Furthermore, the notion of compatibility is extended to include not only pairs of gates, but *n* + 1-Tuples of gates with *n* being the maximum number of inputs of gates in the library (see Methods 4.3). This renders possible determining the exact compatibility of all combinations of gates present in the circuit. Furthermore, it can be asserted that signal levels in a circuit of cascaded compatible gates do not cross the defined thresholds and thus exhibit a defined perturbation margin. By pre-computing the compatibility of gates of a library into an *n* + 1-dimensional matrix, it can be integrated naturally into constructive technology mapping methods like the B&B approach presented in the following.

**Figure 2:**
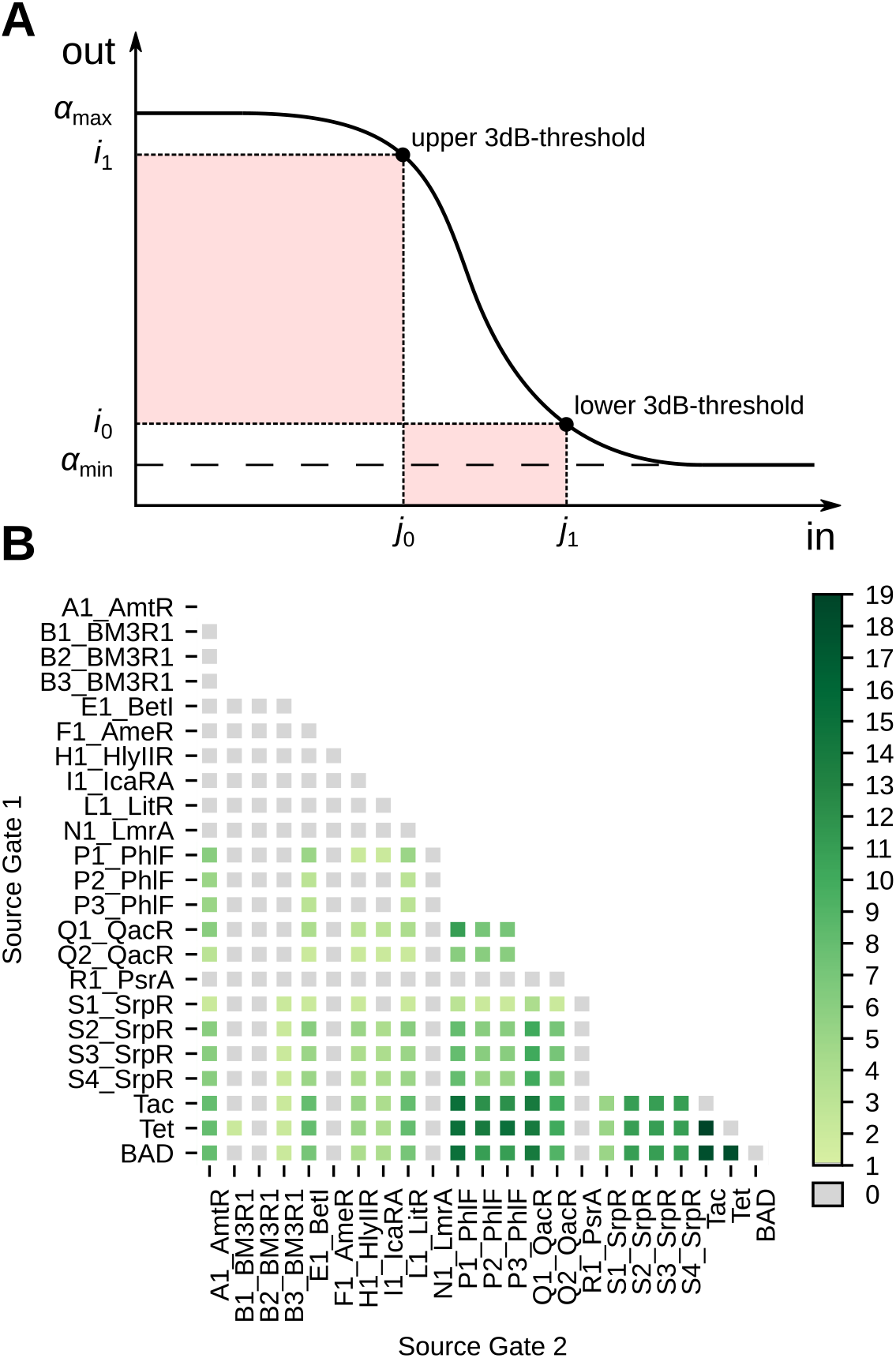
*A*) Visualization of the 3dB-thresholds on which the signal compatibility analysis is based. *i*_0_, *i*_1_ represent the output values at the upper and lower threshold respectively, and *j*, *j*_0_ the corresponding input values. *B*) Compatibility of gates in Cello’s library. For each pair of source gates, the number of compatible target gates is presented. Interestingly, there is a significant number of pairs not compatible with any subsequent NOR gate. This limits the number of gates available for assignment at a distinct position within the circuit.

We have evaluated the compatibility of the NOT and NOR gates from Cello’s library (see Fig. 2). As the library features gates with at most two inputs, the resulting compatibility matrix is 3-dimensional, with Boolean elements representing the compatibility of a triple of two source gates and one target gate. Altogether, 21 % of all triples that can be formed from the library have compatible signal characteristics.

#### 2.3.2 Branch-And-Bound Gate Assignment

The Branch and Bound (B&B) (*27*) method is an optimization strategy for traversing search spaces in an efficient way and obtaining the optimum solution. Generally, in the branching step, partial or complete solutions are generated from other partial solutions according to a branching rule. By this, a search tree is generated iteratively. Then in the bounding phase, the quality of partial solutions is estimated optimistically and compared to the current best known solution. This enables pruning parts of the search tree that spring from inferior partial solutions and thus reaching the optimum more efficiently.

In the context of gate assignment for genetic circuits, the branching rule arises from the fact that the last gate in the cascade defines the maximum output interval of the circuit. Let *x* be the input promoter activity, h(*x*) the inhibitory Hill function modelling the doseresponse curve of the transcriptional regulation, *K* the repression coefficient, *n* the Hill coefficient and [*α*_min_, *α*_max_] the output interval. Then

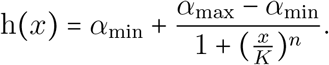

So the circuit’s output interval and thus its maximum score is limited by [*α*_min_, *α*_max_] of the last gate. Upstream gates only determine which portion of the interval is driven by the circuit. Thus, we propose to build partial solutions starting from the output gate and iteratively assign further gates in reversed topological order (see Fig. 3A). This enables pruning considerable parts of the search tree early in the search process. Furthermore, it ensures that every partial solution to the problem represents a valid sub-circuit of the original circuit with new inputs. To estimate the quality of these sub-circuits optimistically, we make use of another feature of the Hill dose-response curve, its monotonicity. If ideal, i.e. maximal input values into the sub-circuit are assumed, it can be stated that these translate to ideal output values and thus score (see Methods 4.4). Thus, by composing the maximally possible input values from the output intervals of the gates left in the library, each partial solution is embedded into an optimistic complete solution (see Fig. 3B). By simulating the sub-circuit configured in this way, the score maximally reachable with a complete solution that contains the partial solution is obtained. Thus, it represents an optimistic estimation of the partial solution’s quality.

**Figure 3:**
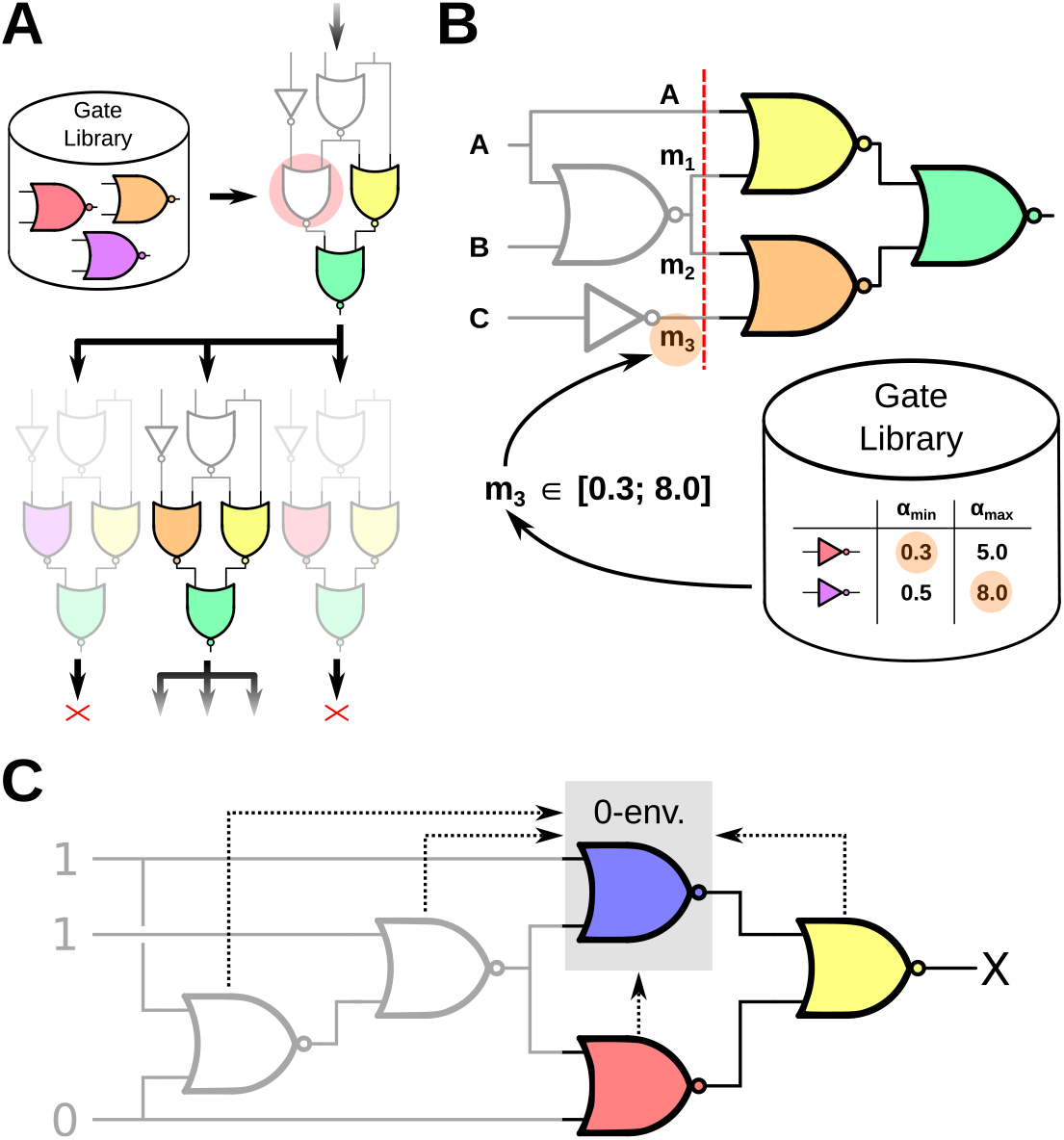
*A*) Excerpt of B&B’s search tree. A branch is performed by mapping gate implementations in the library to the gate marked in red. The formed partial solutions are then scored by the optimistic estimator and pruned, if their estimated quality is below the current best known solution. *B*) Estimation of input intervals exemplary shown for the signal *m*_3_. The optimistic input interval is composed of the parameters [*α*_min_, *α*_max_] of gates left in the library. *C*) Extension of the input interval estimation to crosstalk-environments. The given assignment of Boolean input values leads to an expected “0”-output of the marked gate. Thus, during the estimation of the sub-circuit the gate is embedded into its 0-environment that contains the maximum repressing crosstalk.

The estimation requires valid bounds to the possible signal values to be present at every site of the circuit. That is, every signal representing a Boolean 1 has to be estimated to the highest possible value at this site and vice versa. This requires a further step to be taken at gates with multiple inputs to guarantee optimality. For example, a NOR-gate with two inputs can be decomposed into the superposition of both input values *x* = *x*_1_ + *x*_2_ and the application of the inhibitory Hill function *y* = h(*x*). If input signals of mixed Boolean polarity are assumed, the gate is expected to output a value representing a Boolean 0 by its functional specification. The value of this output signal has to represent a valid lower bound to the signal values possible at this site. Due to the monotonicity of h(*x*), this translates to an upper bound at the gate’s input. However, if both input signals are assumed to be valid bounds, the superposition of these signals does not represent a valid upper bound due to their mixed polarity. Therefore, the output signal of the NOR gate does not represent a valid lower bound. Thus, the input signal representing a Boolean 0 has to be substituted by the highest value possible at this site to guarantee optimality. Generally, this is *α*_max_ of the gate driving the input. This reduces the tightness of the estimation as the substituted value typically deviates strongly from the value present at the site. When mapping a circuit under the constraint of signal-compatibility of gates, io can be used as a substitute instead, as it represents a valid upper bound for the input signal representing a Boolean 0. This restores the tightness of the estimation and thus benefits the efficiency of B&B. Note that the substitution with *i*_0_ only guarantees an optimistic estimation when mapping with compatibility and without crosstalk, as the feedback loops introduced violate the compatibility constraint. Thus, when mapping without compatibility or with crosstalk, we call the substitution with *i*_0_ the “heuristic” mode of B&B, while substituting with *α*_max_ is called “optimal” mode.

So far, the estimation is only applicable to the evaluation of circuits without crosstalk. To find the optimal gate assignment under the influence of crosstalk-effects, it is necessary to include them into the optimistic estimation. To achieve this, we propose the concept of crosstalk-environments in the following. Additionally to the desired inputs of a gate depicted by the wiring diagram, crosstalk introduces further inputs that model the crosstalk effect that other TFs have on the target gate. For each gate, depending on its desired Boolean output according to the functional specification, we compile an optimistic “0-” or “1-environment”. The gate is then embedded into it during the circuit simulation (see Fig. 3C). Besides the optimistic values of the wired inputs, the environments contain the maximum crosstalk-effects that benefit its specified output (see Methods 4.5). In a circuit consisting of gates based on transcriptional repression, e.g., maximum crosstalk from other gates would be assumed in the gate’s 0-environment. In its 1-environment, however, the minimal possible crosstalk is assumed. It is possible to estimate the extremal effect of crosstalk as it is limited by the maximum input values of the gate causing the crosstalk.

The signal-compatibility of genetic gates presented in 2.3.1 can be used to further refine the proposed B&B method as it integrates naturally in the branching step. Naively, the compatibility of all gates in a circuit can be checked as soon as a complete assignment, that is a complete solution, exists. We propose a satisfiability (SAT) based look-ahead approach that is performed after every branching step on the search tree. It checks whether any combination of compatible gates left in the library exists that completes the assignment (see Methods 4.6). By this, the evaluation of sub-trees that lead to invalid full solutions is suppressed as early as possible.

### 2.4 Experimental Results

To evaluate the performance of the proposed gate assignment methods and the behavior of genetic circuits considering crosstalk modeled thermodynamically, we perform the technology mapping for a set of circuits with different parameters. First, we examine the performance of the B&B method in the classic GDA scenario without the consideration of crosstalk. Then, technology mapping is performed with different distributions of crosstalk across the library of genetic gates. This allows conclusions about the performance of B&B and the performance degradation of genetic circuits with crosstalk.

For first evaluating the performance of the proposed B&B method without the consideration of crosstalk, we carried out the gate assignment for the 33 circuit topologies presented in (*23*) using Cello’s library of genetic gates. We then measured the number of circuit simulations needed as well as the deviation from the maximum circuit score, i.e. solution quality, observed in any circuit and compare it to an exhaustive search. Furthermore, all methods have been evaluated both with the application of the compatibility constraint and without.

Table 1 summarizes the results of all gate assignment runs. Without the consideration of compatibility of gates, B&B reduces the number of simulations needed 15-fold compared to exhaustive search, while also guaranteeing to find the optimum solution. In heuristic mode, B&B even exhibits a 664-fold efficiency gain, while the worst solution quality observed deviates only −2.7 × 10^-3^ % from the maximum. The application of the compatibility constraint reduces the search space to 1,25% of the original one. In this reduced space, the optimal B&B exhibits a 108-fold gain in the number of needed simulations compared to exhaustive search. Finally, the gate assignment has also been carried out with Cello’s stochastic simulated annealing (SA) optimization method. Note the limitation that Cello applies a similar but not identical notion of compatibility (see 2.3.1). Furthermore, due to the stochastic nature of SA, all runs have been carried out 10 times and the worst case deviation has been determined by comparing the best and the worst solution quality observed for each circuit. Cello’s SA features a fixed number of 30.000 simulations per circuit, leading to a total number of 990.000 simulations needed, while a worst case deviation of −65 % has been observed. B&B in comparison finds the optimum solution deterministically and exhibits a 83-fold efficiency gain.

**Table 1:**
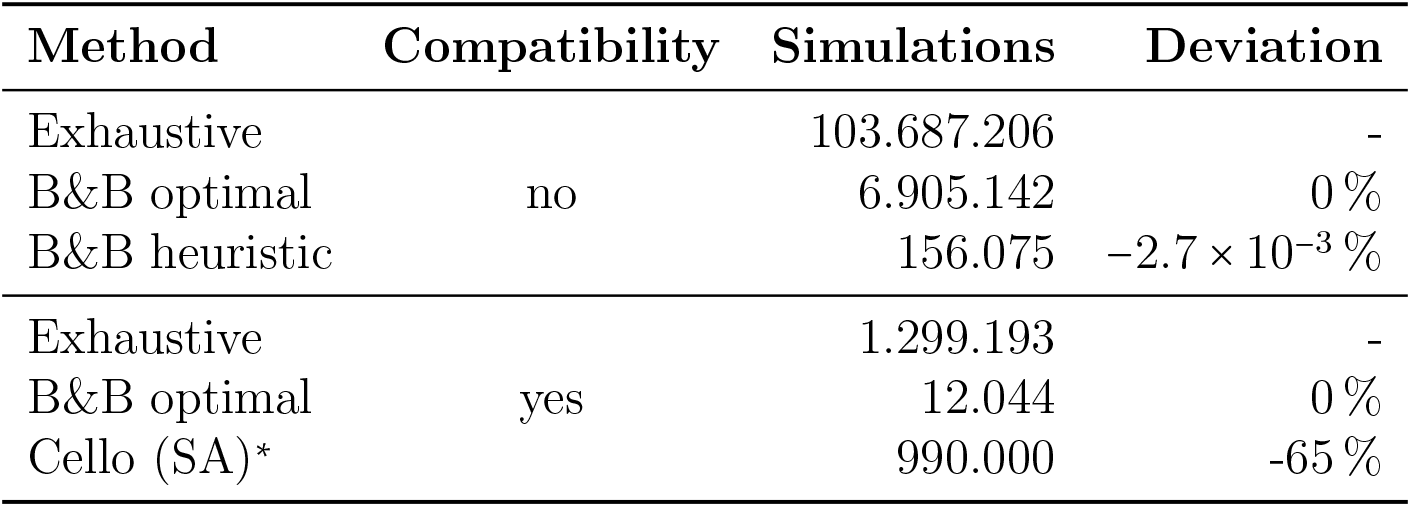
Number of circuit simulations needed and worst case deviation of the reached circuit score for mapping the set off 33 circuits presented in (*23*) without consideration of crosstalk. * Cello uses a similar, but not identical notion of compatibility in which approx. 30 % instead of 21 % of gate triples are considered compatible.

| Method | Compatibility | Simulations | Deviation |
| --- | --- | --- | --- |
| Exhaustive | no | 103.687.206 | - |
| B&B optimal |  | 6.905.142 | 0 % |
| B&B heuristic | | 156.075 | $-2.7 \times 10^{-3}$ % |
| Exhaustive | yes | 1.299.193 | - |
| B&B optimal |  | 12.044 | 0 % |
| Cello (SA)* |  | 990.000 | -65 % |

Figure 4F relates the mean number of simulations needed per circuit and mapping method to the circuit size. The exponential problem complexity when not considering gate compatibility can clearly be seen. However, compared to exhaustive search, optimal B&B reduces the number of simulations by up to 2.4 orders of magnitude and heuristic B&B again by up to 1.6 orders of magnitude. When considering compatibility, the problem complexity is diminished due to the number of allowed gate assignments decreasing with increasing circuit size. In this case, the refined estimation of partial solutions enables optimal B&B to reduce the number of simulations up to 2.6 orders of magnitude,

**Figure 4:**
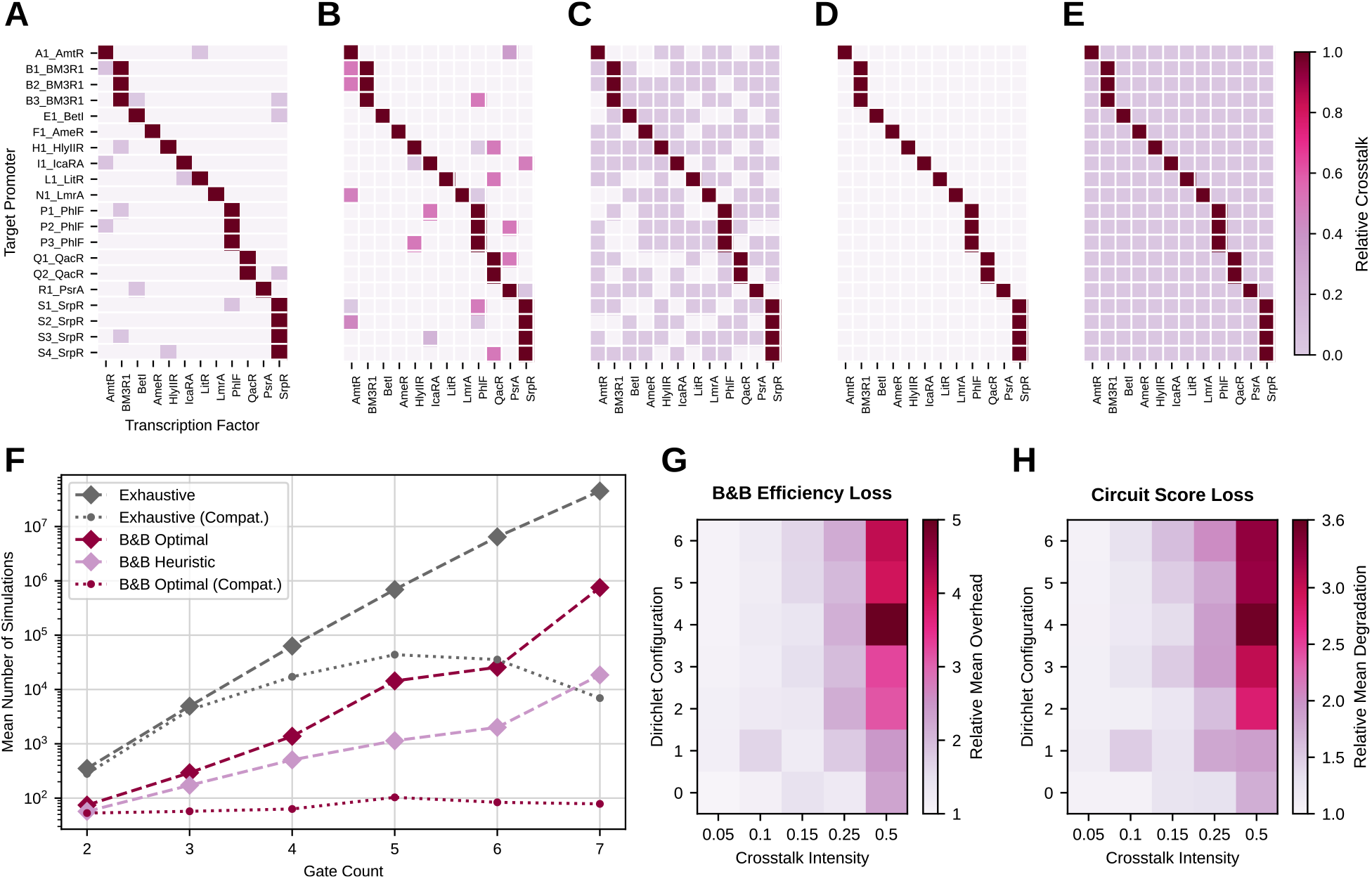
*A*) – *E*) Different distributions and total intensities of the considered crosstalk. A) very peaked distribution, low total intensity; *B*) very peaked distribution, large total intensity; *C*) distribution with moderate entropy, moderate total intensity; *D*) distribution almost uniform, high entropy, low total intensity; E) distribution almost uniform, high entropy, large total intensity. *F*) Mean number of simulations needed for mapping circuits with the proposed B&B methods compared to exhaustive search with respect to the number of gates. All results in this plot depict technology mapping runs without crosstalk. Dashed lines show results without the compatibility constraint, while dotted lines depict results with compatibility. *G*) Number of simulations needed for mapping the set of 66 circuits with different crosstalk configurations in relation to the result without crosstalk. As a mapping method, optimal B&B with consideration of compatibility has been used. *H*) Mean score of the 66 circuits mapped with optimal B&B with different crosstalk configurations in relation to the scores reached without crosstalk. The color encodes the multiplier needed to map the score with crosstalk to the one without crosstalk.

In the following, we evaluate the influence of crosstalk on the performance of the B&B gate assignment and genetic circuits by performing technology mapping with multiple gate libraries containing different crosstalk configurations. The crosstalk configurations were generated in the following way. First, for each available promoter with index *m*, a random vector of values 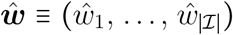, one for each non-cognate TF *k* ≠ *m*, was generated by sampling a Dirichlet distribution with a single concentration hyperparameter *γ* > 0. This allowed us to control how sparse the random crosstalk will be distributed across the noncognate TFs. These values were then used to obtain normalized relative energy terms of the binding state of the cognate TF at the promoter 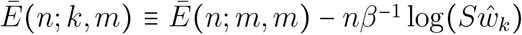 for 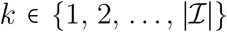 and a number *S* > 0. The number *S* poses as a total intensity hyperparameter. This ensured that at 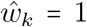 and *S* = 1, a non-cognate TF would repress transcription at the promoter equivalently to the cognate TF. Since the entries of any sample from a Dirichlet distribution are guaranteed to sum to one, i.e. 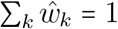, introduction of *S* allowed us to control the total crosstalk coming from all non-cognate TFs. We generated 35 libraries for each combination of one of 7 different concentrations *γ* and one of 5 total intensities *S*. The *γ* and *S* used are given in the supplementary material but five libraries are visualized in Fig. 4A-E. Additionally, the set of 33 circuits is extended with structural variants synthesized in (*19*) to form a set of 66 circuits. We could observe that for low to moderate total intensity in crosstalk only a small deterioration in either score of the found assignment or number of simulations w.r.t. the crosstalk-free case occurred. From around 15 % total intensity on, the excess number of simulations and the reduction in score depended strongly on the concentration hyperparameter as shown in Fig. 4G-H. For very peaked crosstalk distributions, the assignments were found in comparable time to the case of low intensity, but their scores were at around 50 % to 60 % of the crosstalk-free case. In this case, the acceptable results can be achieved by either distributing crosstalk across the gates to match their desired logic outputs or by mitigating assignments with gate combinations exhibiting crosstalk at all. For crosstalk distributions with high entropy on the other hand, the strongest observed decrease in score and the largest excess in number of simulations has been observed. In the most extreme case, we observed a reduction to 20 % in average score and an 5-fold increase in number of simulations. In this scenario, with rising total intensity it seems to be increasingly hard to find assignments that correctly implement the logic function. Also the B&B algorithm’s bounding function becomes increasingly non-descriptive w.r.t. prediction of the final score which leads to the its search strategy starting to approach that of an exhaustive search across the assignments. In the observed worst-case crosstalk configuration, the heuristic B&B method exhibits a 10.5-fold efficiency gain compared to the optimal method, while obtaining near optimal results that feature a mean −0.7% deviation from the optimum score. Thus, heuristic B&B represents a viable alternative for efficient technology mapping of circuits that are subject to crosstalk. The non-smoothness of the color plots in Fig. 4G-H can be explained by the fact that each library represents a single sample of a random library with fixed total intensity and concentration.

## 3 Conclusion

In this contribution, we presented a novel technology mapping approach for GDA which can process device libraries that exhibit crosstalk between different transcription factors. This crosstalk leads to binding of TFs at non-cognate binding sites and thus leads to unintended repression or activation. The approach is also capable of dealing with titration effects which reduces the number of TFs available for binding at the cognate binding sites due to off-target site on the host genome. Both effects have to be taken into account during circuit scoring in order to find robust designs for genetic circuits, in particular, for combinational logic circuits. The underlying principle of our developed circuit models uses thermodynamic calculations of TF-DNA binding energies. We have shown that using this modeling approach we can find good gate assignments even in the presence of both context effects.

We investigated different types of crosstalk scenarios. They vary in the intensity and in the entropy of the crosstalk distribution, i.e., whether crosstalk is spread equally among the library parts or whether crosstalk is confined to a few pairs of parts of the library. It turns out that a certain amount of crosstalk can be tolerated in the library with a moderate degradation in circuit performance, quantified in terms of circuit score. For higher crosstalk levels, the circuit performance degrades significantly and the computational complexity of the proposed B&B assignment method increases concurrently. In case of severe crosstalk but with low entropy, the B&B run-time showed almost no degradation, but the circuit performance was strongly affected. In worst-case crosstalk scenarios, the heuristic B&B assignment still found near optimal results, yet still 13 times faster than the optimal, exhaustive method.

We believe that these findings will help other researchers to evaluate their device libraries at an early stage during research and circuit development. They can judge the crosstalk situation in their libraries and in their circuit designs without the need to build many different test circuits in the lab.

## 4 Methods

### 4.1 Thermodynamic Description of the Genetic Logic Circuit

In the following sections, we derive an equation which allows to express the TF concentration *f_m_* of a gate with index 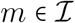 as a function of all TF concentrations *f_k_*, 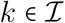 of the other gates and any additional nuisance TF’s in the circuit if present. Since the circuit is comprised of NOR gates that map promoter activities to promoter activites, we then reformulate the expression to obtain the transfer functions of the gates (5) and as a simplification (3).

#### 4.1.1 Gate-Internal TF Expression

We first present the general expression for the partition function of bound RNAP for the NOT (1-input NOR) gate’s output promoter under potential crosstalk from all *K* available TF in the circuit. The quantity *p* is the number of available RNAP.

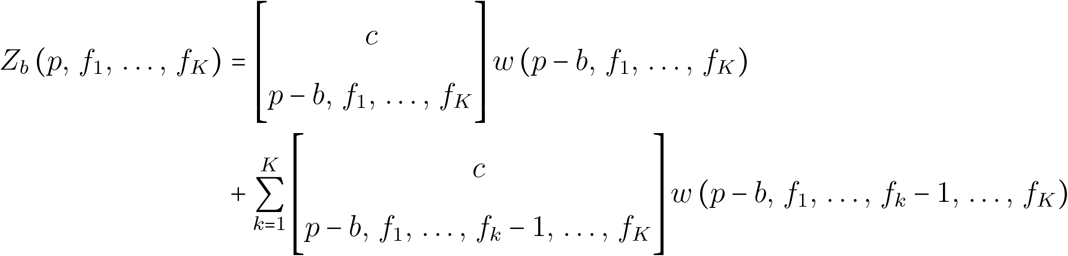

with *b* ∈ {0, 1}, the completed multinomial coefficient

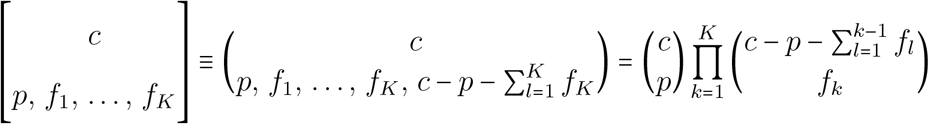

and the general statistical weight function *w*

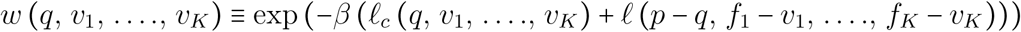

which covers the partition across the background and specific binding sites. For the background energy we assume independent binding. Thus, the function *£_c_* takes on the simple form

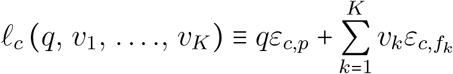

with the absolute binding energies *ε_c,p_* for RNAP and *ε_c,f_k__* for TF binding to a non-specific background site. If the specific binding energy function *ℓ* can be written as a linear combination of only a few energy terms for most arguments depends on the degree of simplification in the model. A fully independent specific binding would lead to a similar appearance to ℓ_*c*_. The particular choice for ℓ implements e.g. cooperativity and basal expression. Generally, we obtain the expression for the expectation E (*X* | *f*_1_,…, *f_K_*) via

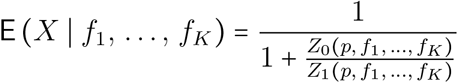

The derivation of the final simplified expression is takes a lot of space, so we refer to the supplementary material for it. Under the assumption of no relevant RNAP titration, i.e. a sufficiently large copy number of RNAP’s available for binding, and no cooperativity, we then obtain as the result for the NOT gate with imperfect competition and arbitrary crosstalk from other TF’s

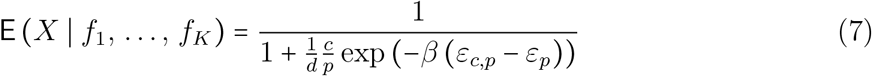

where the regulation factor *d* is given by

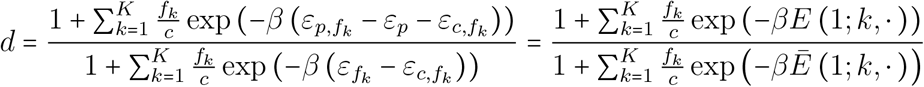

where we introduced the relative energy functions *E* (1; *k, ·*) ≡ *ε_p_,f_k_* - *ε_p_* - *ε_c,f_k__* and 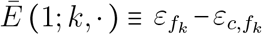 which encode the energy expense of the *k*-th TF binding to the promoter which has no index and thus obtains a · here. Note, that *d* in the NOT case is approximately equal to the fold-change *ϕ* (*f*_1_, …, *f_k_*). To extend the result to the cooperative case, we can simply follow the construction for dependent multiple bindings from (*21*).

In the following, we will add the superscript (*n*) to any “anonymous” input promoter for enumeration. In this way, there is no confusion with the gate index placed as a subscript and simultaneously allows us to specify an arbitrary *n*-th input promoter.

The output expression in the N-input NOR gate is given by a simple sum of the expressions for NOT gates with the same output gene. This means that the decomposition in the single expectations 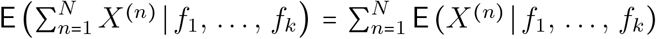 yields a sum of expressions that match those of the NOT case. For this to happen, the random variables *X*^(*n*)^ need to be independent. We can argue analogous to (*25*) that they are independent under the assumption of no relevant intra-circuit inter-promoter TF and RNAP titration. This means that, conditioned on the binding of a TF or RNAP to binding sites of another gate, the probability of binding to the sites of the considered gate doesn’t significantly change. This can be seen as a “large” copy-number approximation, while “large” will be relative to the potential binding sites present in the circuit.

It is now a straight forward task to construct the expression one for the *n*-input NOR gate. Thus, in this case we directly obtain

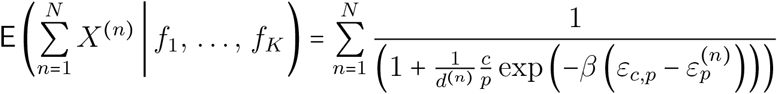

with the promoter-specific regulation factors *d*^(*n*)^ given by

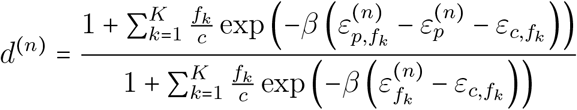

which is the final expression for the output expression of the (*N*-input) NOR gate with imperfect competitivity (between RNAP and TF’s) and crosstalk from an arbitrary number of additional TF’s.

This is the first case, where we are now interested in an expression for the overall fold-change. The fold-change is given by the expression

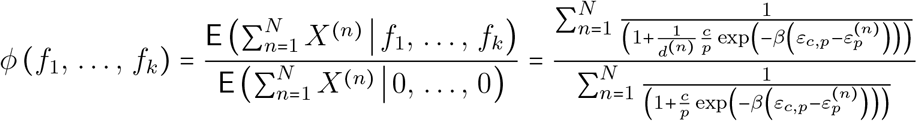

which we can approximate closely by assuming, that the factors 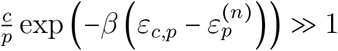 for all *n*. This is based on the assumption, that the expectation of encountering an RNAP bound at the specific promoter is in absolute numbers rather small, i.e. Pr (*X*^(*n*)^ = 1|*f*_1_, …, *f_k_*) is small for all *n* and the number of binding sites is reasonably small as well (it is even only one here at this moment). Also *d*^(*n*)^ ≤ 1 holds true anyway and even amplifies this aspect for the scaled terms 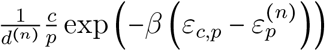. Then, we get for the fold-change

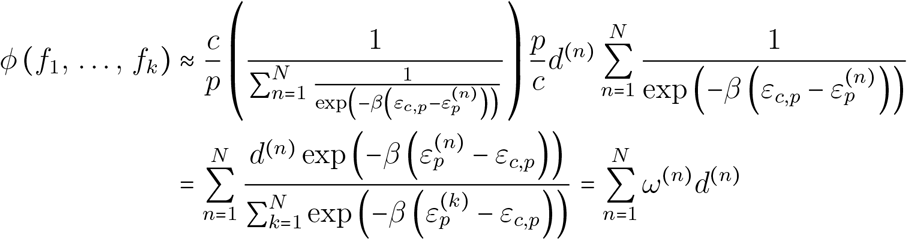

which lets us state the final form for the fold-change by

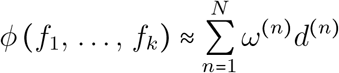

where the weights *ω*^(*n*)^ are given by the equation

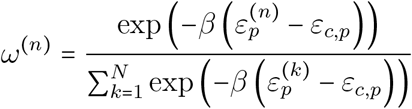

which relates the relative statistical weights of the promoters to each other. Finally, we repeat the expression for the regulation factors *d*^(*n*)^, which is given by

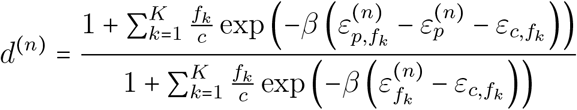

We again refer to (*21*) on how to extend the *d*^(*n*)^ to the case with cooperativity. Actively repressing stronger promoters has a stronger influence on the overall fold-change of the NOR gate. The fold-change together with a proportionality constant then gives the absolute expression level *f_m_*, i.e.

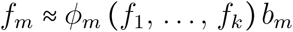

for some *b_m_* ∈ ℝ_≥0_. In accordance to previous works on thermodynamic gene expression (*21*), since *ϕ_m_* ∈ (0, 1] and argmax_*f*_1_…*f_K_*_ *ϕ_m_* (*f*_1_, …, *f_K_*) = (0,…,0), the proportionality constant *b_m_* = max *f_m_* equals the unrepressed, maximal TF concentration.

#### 4.1.2 From TF Concentrations to Promoter Activity

While being intuitive, working with mappings of TF concentrations raises consistency issues with the gate and circuit models. This stems from the fact, that TF concentrations are gate-internal quantities and thus any change in wiring would change the “transfer” functions to compute, This is also incompatible with common methods to improve performance, e.g. lookup tables of function values. Thus, to keep logic circuit and thermodynamic models consistent, we need to formulate the output promoter activities *α_m_* as functions of the other promoter activities *α*_1_,…,*α_K_*. First, we assume the output promoter activities *α_m_* ∝ E (*X_m_* |*f*_1_,…,*f_K_*) to be proportional to their occupancies by RNAP. Thus, we obtain for the output activity that *α_m_* ≈ *ϕ_m_* (*f*_1_,…, *f_k_*) *q_m_* with a new proportionality constant *q_m_* ∈ ℝ_≥0_. Clearly, by the same argument as used for *b_m_*, we can see that *q_m_* = max*α_m_*, i.e. *q_m_* must be the maximum unrepressed promoter activity. Thus, we obtain for the relationship between input promoter activity and TF concentration at the *m*-th gate

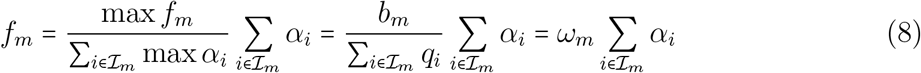

The final step is to relate the internal TF concentration *f_m_* and the output promoter activity *α_m_*. Since we decomposed the NOR gate’s input to a sum of inputs of NOT gates which is already reflected by (8), it is sufficient to consider this relation w.r.t. the fold-change of a NOT gate. Then, we obtain from (7) that

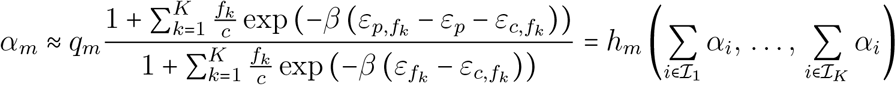

### 4.2 Iterative Calculation of the Circuit Response

Before we calculate the circuit response under crosstalk (6), we start by considering the crosstalk-free version (4). Calculation of (4) is very fast because there exists at least one sorting of the 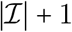 equations in the system, such that each equation can be solved explicitly upon solution of the previous one. As a consequence, a single iteration, i.e. a single calculation of each equation in the system, is sufficient to obtain the final solution *y* given 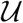. Thus, under the assumption that in most assignments traversed by the technology mapping process the incorporation of crosstalk through (6) does not fundamentally change the gate’s outputs in comparison to (4), we use the solution of (4) as an initial guess. As a nonlinear equation system, we can write (6) in matrix form with the vector of unknowns 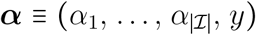 including the response *y*. Then, with the fixed vector-valued function 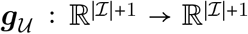 which maps gate-outputs to gate-outputs, we can solve the fixedpoint problem **0** = ***α*** – ***g*** (***α***) with the vector of zeros **0**. This can then together with the initial guess ***α***_0_ obtained from (4) be fed into a state-of-the-art fixedpoint or root solver. For our evaluation, we simply used Python’s `scipy.optimize` package with the function root and chose the Levenberg-Marquardt algorithm as a solution method.

### 4.3 Defining Signal-Compatibility of Genetic Logic Gates

Let [*α*_min_, *α*_max_] be the output interval of a genetic gate. Then 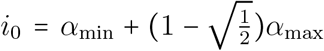 represents its output at the lower 3dB-threshold and *j*_1_ = g^-1^(*i*_0_) the corresponding input value. Similarly, 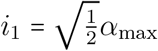 is the output at the upper 3dB-threshold and *j*_0_ = g^-1^(*i*_1_) its corresponding input value.

Assume a library of gates based on repression with a maximum of two inputs per gate. A triple of two source gates *r, s* and a target gate *t* is compatible if

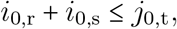

i.e., the superposition of the maximum input values representing a Boolean 0 does not exceed the lower input threshold of the target gate. Further,

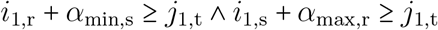

ensures that the lowest input superposition representing a Boolean 0 and a Boolean 1 does not fall below the target’s upper input threshold.

### 4.4 Optimality of the Branch and Bound Method

Let *γ* be the topology of the genetic circuit and *a* the assignment of gates from the library to the abstract nodes of the circuit. Let further be 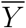 the set of output signal values representing a Boolean 1 and 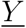 the signal values representing a Boolean 0. Then the circuit score (9a) introduced by Cello rates the minimum separation of the complementary Boolean output states of the final gate in the circuit (*23*) (*28*).

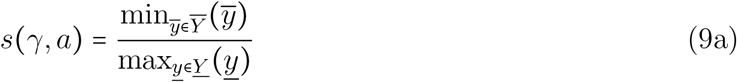

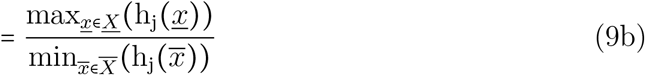

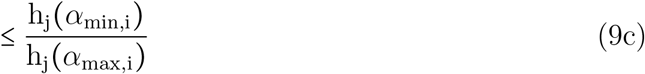

Let *j* name that final circuit gate and let h_j_(*x*) be its transfer function, the inhibitory Hill curve. Let 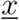 be the maximum gate input value of all input values 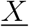 representing a Boolean 0, 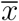 be the minimum gate input value of all input values 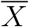 representing a Boolean 1. Then (9b) results from the monotonicity and inverting characteristic of h_j_(*x*). Let [*α*_min;i_, *α*_max;i_] be the bounded output interval of the preceding gate *i*, then (9c) represents a valid upper bound of the score, because 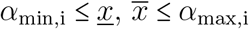.

### 4.5 Compilation of the Crosstalk-Environments

Let *g* ∈ *G* be the gate in the set of gates *G* comprising the abstract circuit topology for which the crosstalk-environment shall be built. Assume that it is assigned with a genetic gate implementation *t* ∈ *D* from the set *D* of implementations in the library. The crosstalk-environment of *g* is built by superimposing the contributions of all other gates *h* ≠ *g* ∈ *G* in the circuit. Let furthermore [*c*_min,h_, *c*_max,h_] be the minimum and maximum input values into source gate *h* determining its output and thus crosstalk induced by it. If the input gates of *h* are assigned an implementation from the library, this interval comprises the superposition of the input gates’ minimum and maximum output values respectively. If the input gates of *h* are unassigned, the interval is composed of the minimum and maximum output values of gates left in the library, representing an extremal estimation.

First, assume that the source gate *h* is assigned an implementation *s* ∈ *D* and its cognate TF is *b_s_*. Its contribution to the crosstalk-environment of gate *g* is then calculated according to the input values and expressed TFs shown in Table 2. To guarantee an optimistic estimation of the partially assigned circuit, its produced crosstalk is assumed to be maximal in the target gate’s 0-environment and minimal in the 1-environment. As the source gate’s TF is fixed, this is done by minimizing or maximizing its possible input values.

**Table 2:** Source gate activity and TF assumed in crosstalk-environments if the source gate *h* is assigned an implementation *s* ∈ *D* with cognate TF *b_s_* for optimistic estimation of crosstalk.

| | Source Gate ($s \in D$) | | |
| --- | --- | --- | --- |
|  | env. | 0 | 1 |
| Target Gate ($t \in D$) | 0 | $c_{\max,h}, b_s$ | $c_{\max,h}, b_s$ |
| | 1 | $c_{\min,h}, b_s$ | $c_{\min,h}, b_s$ |

Let *b*_min,t_ and *b*_max,t_ be the non-cognate TFs of gate implementation *t* assigned to target gate *g* that are not contained in the partial assignment and have the minimum and maximum binding affinity to *t*’s promoter respectively. The contribution to the crosstalk of a source gate that is not assigned is then determined according to Table 3. That is, additionally to assuming extremal input values for the crosstalk-inducing gate, it is assumed to express the TFs that have the minimum or maximum effect on the target gate.

**Table 3:**
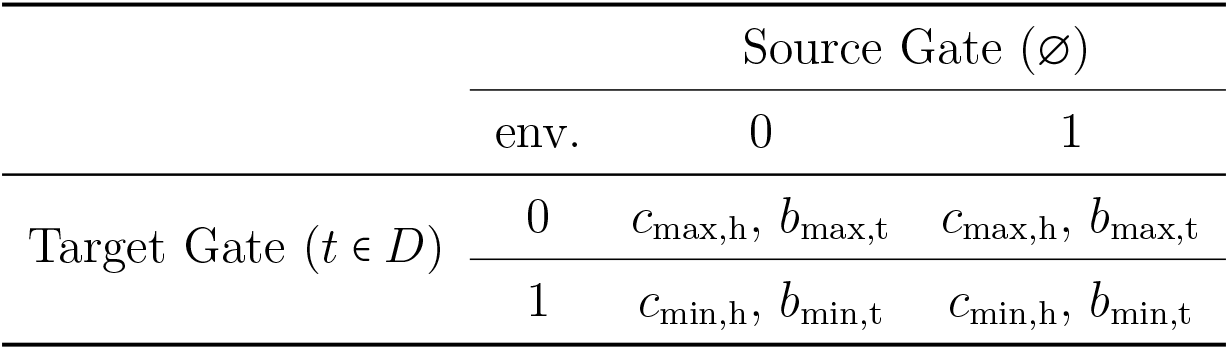
Source gate activity and TF assumed in crosstalk-environments if the source gate *h* is not assigned for optimistic estimation of crosstalk.

| | Source Gate (∅) | | |
| --- | --- | --- | --- |
|  | env. | 0 | 1 |
| Target Gate ($t \in D$) | 0 | $c_{\max,h}, b_{\max,t}$ | $c_{\max,h}, b_{\max,t}$ |
| | 1 | $c_{\min,h}, b_{\min,t}$ | $c_{\min,h}, b_{\min,t}$ |

The choice of the composition of the crosstalk-environments depend only on the specified Boolean output value of the target gate. Furthermore, feedback loops introduced by crosstalk are severed by basing c_min_,_h_ and c_max_,_h_ solely on minimum and maximum activity values given in the library which represent physical bounds. Both aspects ensure that the estimation of the crosstalk is fully optimistic w.r.t. to the functional specification.

Analog to the heuristic mode introduced in 2.3.2 for the estimation of wired inputs, we can estimate the crosstalk heuristically. Let *j*_0,h_ and *j*_1;h_ be the input values of source gate h corresponding to the thresholds introduced in the signal-compatibility analysis 2.3.1. For circuits built from compatible gates, we can state that *j*_0,h_ represents a valid upper bound for the possible inputs of gate *h* when its specified output is a Boolean 1. Analog, *j*_1;h_ represents a valid lower bound for the possible inputs when the specified output of gate h is a Boolean 0. Note that these bounds are only valid for a circuit free of crosstalk, i.e. without feedback loops. Nonetheless, they can be used as a heuristic estimation for the output of h when considering circuits with crosstalk. We implemented this estimation into the heuristic mode of B&B according to Tables 4 and 5. In contrast to the optimal mode, the optimistic but unrealistic estimation of the input activity *c*_max,h_ in the case of a specified Boolean 1 output of gate *h* is replaced by *j*_0,h_. Analog, *c*_min,h_ is replaced by *j*_1;h_ in the case of a Boolean 0 output.

**Table 4:** Source gate activity and TF assumed in crosstalk-environments if the source gate *h* is assigned an implementation s ∈ *D* with cognate TF *b_s_* for heuristic estimation of crosstalk.

| | Source Gate ($s \in D$) | | |
| --- | --- | --- | --- |
|  | env. | 0 | 1 |
| Target Gate ($t \in D$) | 0 | $c_{\max,h}, b_s$ | $j_{0,h}, b_s$ |
| | 1 | $j_{1,h}, b_s$ | $c_{\min,h}, b_s$ |

**Table 5:** Source gate activity and TF assumed in crosstalk-environments if the source gate h is not assigned for heuristic estimation of crosstalk.

| | Source Gate (∅) | | |
| --- | --- | --- | --- |
|  | env. | 0 | 1 |
| Target Gate ($t \in D$) | 0 | $c_{\max,h}, b_{\max,t}$ | $j_{0,h}, b_{\max,t}$ |
| | 1 | $j_{1,h}, b_{\min,t}$ | $c_{\min,h}, b_{\min,t}$ |

### 4.6 Branching with Compatibility Look-Ahead

After every branching step during the B&B, a look-ahead check is performed whether the formed partial assignment of genetic gates can be completed to form a valid solution that meets the constraints given by the compatibility matrix. To this end, a satifiability (SAT) problem is formulated that is solved by a SAT solver online during technology mapping. The problem consists of four clauses, one of which is optional. Let *g* ∈ *G* be a gate *g* from the set of all abstract gates in the circuit *G* and *d* ∈ *R* ⊂ *D* the genetic gate implementation *d* from repressor group *R* that forms a part of all implementations *D* present in the library. Then 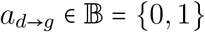 is the Boolean variable representing an assignment of implementation d to gate *g*. Let further be *y* ∈ *Y* the logic type *y* of all logic types present in the library and t(*g*): *G* → *T*, t(*d*) : *D* → *T* the functions mapping gates and implementations to their logic type. Then the first clause of the SAT problem

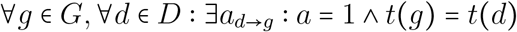

ensures that every gate is assigned at least one implementation of the same type. Further, the clause

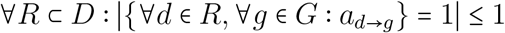

states that every repressor must be assigned maximally once. Let furthermore be *n* the maximum number of inputs of gates in the library and (*s*_1_, *s*_2_… *s_n_,t*) ∈ *T* the *n* + 1-tuple of *n* source implementations *s_n_* and target implementation *t* of all *n* + 1-tuples *T* present in the circuit. Let 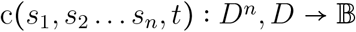 be the function that queries the compatibility matrix for a given *n* + 1-tuple of implementations. Then the clause

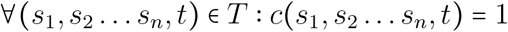

ensures that every tuple present in the circuit consists of compatible implementations. The final clause

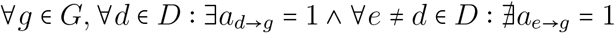

states that every gate is assigned maximally one implementation. It is optional for evaluating the compatibility, but can be useful as it enables the retrieval of valid assignments from the SAT model. To speed up the evaluation of the SAT problem, it is set up prior to the technology mapping process based on the circuit structure and the library of gate implementations. When a (partial) assignment shall be evaluated, the variables *a* are substituted with constants according to the existing assignment.

The source code of the proposed methods is available at https://www.rs.tu-darmstadt.de/ARCTIC.

## 5 Acknowledgements

Nicolai Engelmann and Heinz Koeppl acknowledge support from the European Research Council (ERC) within the CONSYN project, grant agreement number 773196.

## 6 Supporting Information

## 7 Author Contribution

H.K. and C.H. provided the research idea and contributed methodology. N.E., T.S. and E.K. conceived novel modelling and technology mapping schemes and carried out mathematical analysis and software development. All authors contributed to the writing of the paper.

## Graphical TOC Entry

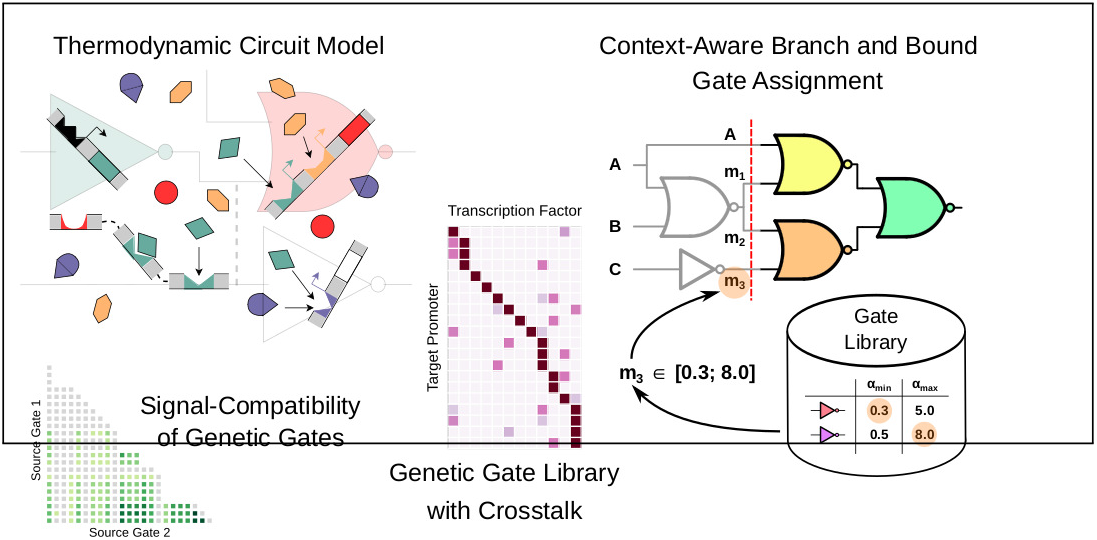

