## Supporting Information for "Context-Aware Technology Mapping in Genetic Design Automation"


Nicolai Engelmann,[†,¶] Tobias Schwarz,[†,¶] Erik Kubaczka,[†,¶] Christian Hochberger,[†,‡] and Heinz Koeppl[*,†,‡]

*†Department of Electrical Engineering and Information Technology, TU Darmstadt, Darmstadt, 64283, Germany*

*‡Centre for Synthetic Biology, TU Darmstadt, Darmstadt, 64283, Germany*

*¶The authors contributed equally to this research.*


#### A Comparison of Thermodynamic Library to Original Cello Library

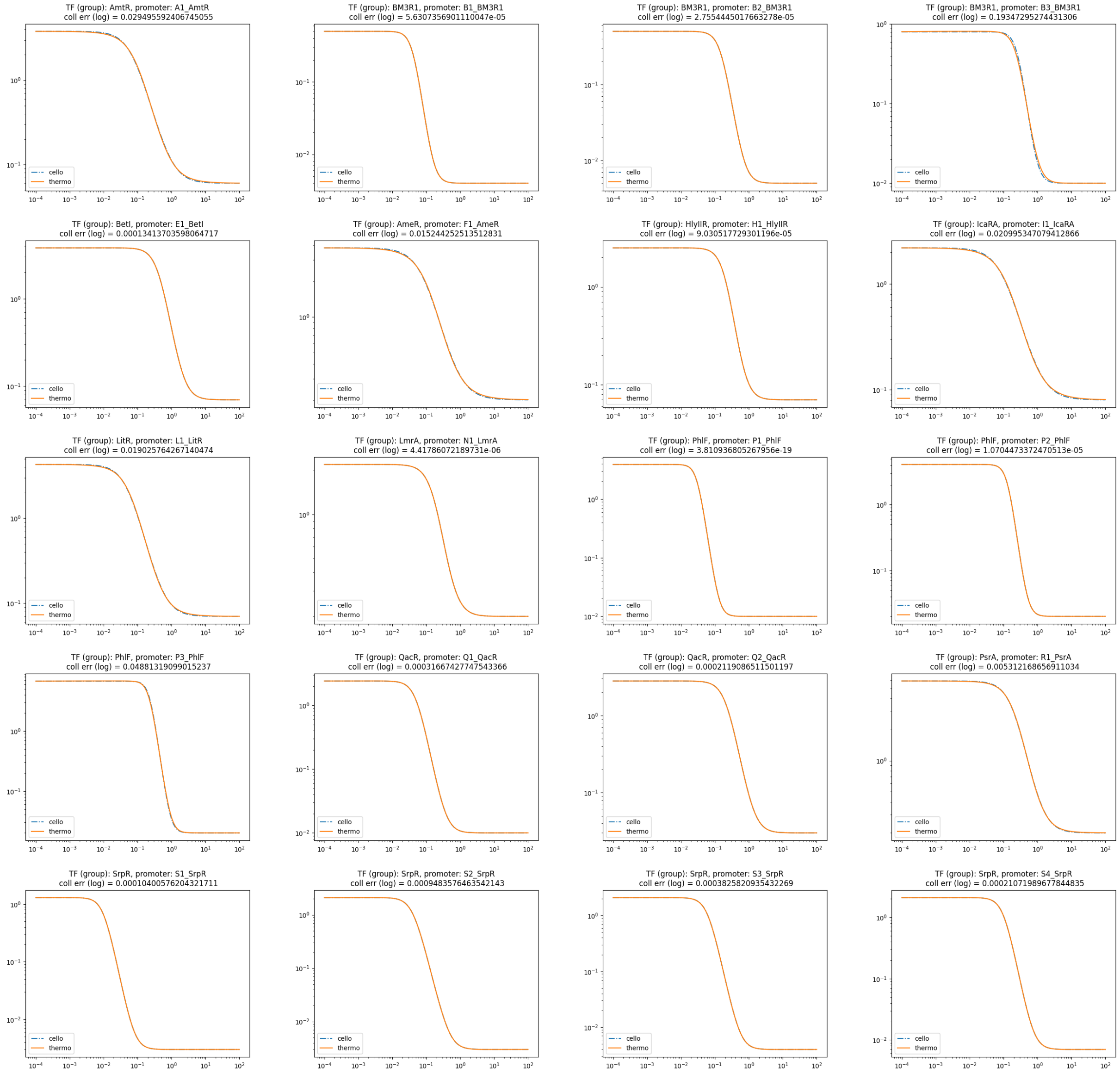


Figure 1: Plots comparing the original Cello transfer functions and the calibrated thermodynamic transfer functions.

#### B Proof of Transfer Function under Crosstalk

We start out with the equation from the main text

$$\mathsf{E}\left(X \mid f_1, \ldots, f_K\right) = \frac{1}{1 + \frac{Z_0(p, f_1, \ldots, f_K)}{Z_1(p, f_1, \ldots, f_K)}}$$

What we did avoid before, we have to tackle now: the case, where we build the ratio of two implicitly completed multinomial coefficients with two arbitrary decrements. We thus seek to simplify

$$\begin{aligned}
&\left[\begin{matrix} c \\ a_1, \ldots, a_K \end{matrix}\right] \Bigg/ \left[\begin{matrix} c \\ a_1, \ldots, a_m - 1, \ldots, a_n - 1, \ldots, a_K \end{matrix}\right] \\
&= \prod_{k=1}^{K} \binom{c - \sum_{l=1}^{k-1} a_l}{a_k} \\
&\quad \Bigg/ \left( \prod_{k=1}^{m-1} \binom{c - \sum_{l=1}^{k-1} a_l}{a_k} \binom{c - \sum_{l=1}^{m-1} a_l}{a_m - 1} \prod_{k=m+1}^{n-1} \binom{c - \sum_{l=1}^{k-1} a_l + 1}{a_k} \binom{c - \sum_{l=1}^{n-1} a_l + 1}{a_n - 1} \prod_{k=n+1}^{K} \binom{c - \sum_{l=1}^{k-1} a_l + 2}{a_k} \right) \\
&\approx \frac{c - \sum_{l=1}^{m-1} a_l}{a_m} \prod_{k=m+1}^{n-1} \frac{c - \sum_{l=1}^{k} a_l}{c - \sum_{l=1}^{k-1} a_l} \\
&\quad \times \left( \binom{c - \sum_{l=1}^{n-1} a_l}{a_n} \prod_{k=n+1}^{K} \binom{c - \sum_{l=1}^{k-1} a_l}{a_k} \Bigg/ \binom{c - \sum_{l=1}^{n-1} a_l + 1}{a_n - 1} \prod_{k=n+1}^{K} \binom{c - \sum_{l=1}^{k-1} a_l + 2}{a_k} \right)
\end{aligned}$$

For the second line to simplify, we need to take a look at a combination of both formulas used before, i.e.

$$\binom{n-k+1}{m-1} \approx \frac{m}{n-k+1} \binom{n-k+1}{m} \approx \frac{m}{n-k+1} \frac{n-k}{n-k-m} \binom{n-k}{m} \approx \frac{m}{n-k-m} \binom{n-k}{m}$$

and thus

$$\binom{c - \sum_{l=1}^{n-1} a_l}{a_n} \Bigg/ \binom{c - \sum_{l=1}^{n-1} a_l + 1}{a_n - 1} \approx \frac{c - \sum_{l=1}^{n} a_l}{a_n}$$

The only thing that's left to derive is an expression for the two-times-increment. For this, we need to apply one of the formulas twice, i.e.

$$\binom{n-k+2}{m} \approx \frac{n-k+1}{n-k-m+1}\binom{n-k+1}{m} \approx \frac{n-k+1}{n-k-m+1}\frac{n-k}{n-k-m}\binom{n-k}{m} \approx \left(\frac{n-k}{n-k-m}\right)^2\binom{n-k}{m}$$

and thus

$$\binom{c-\sum_{l=1}^{k-1} a_l}{a_k} \Bigg/ \binom{c-\sum_{l=1}^{k-1} a_l + 2}{a_k} \approx \left(\frac{c-\sum_{l=1}^{k} a_l}{c-\sum_{l=1}^{k-1} a_l}\right)^2$$

This together then allows us to state

$$\begin{bmatrix} c \\ a_1, \ldots, a_K \end{bmatrix} \Bigg/ \begin{bmatrix} c \\ a_1, \ldots, a_m - 1, \ldots, a_n - 1, \ldots, a_K \end{bmatrix}$$
$$\approx \frac{c-\sum_{l=1}^{m-1} a_l}{a_m} \prod_{k=m+1}^{n-1} \frac{c-\sum_{l=1}^{k} a_l}{c-\sum_{l=1}^{k-1} a_l} \frac{c-\sum_{l=1}^{n} a_l}{a_n} \prod_{k=n+1}^{K} \left(\frac{c-\sum_{l=1}^{k} a_l}{c-\sum_{l=1}^{k-1} a_l}\right)^2$$

and finally carry out the simplification of the expression for the expectation $\mathsf{E}\left(X \mid f_1, \ldots, f_K\right)$, giving us

$$\begin{aligned}
&\frac{Z_0\left(p, f_1, \ldots, f_K\right)}{Z_1\left(p, f_1, \ldots, f_K\right)} \\
&\approx \frac{w\left(p, f_1, \ldots, f_K\right) \quad \cdots}{\frac{p}{c}\prod_{n=1}^{K} \frac{c-p-\sum_{l=1}^{n-1} f_l}{c-p-\sum_{l=1}^{n} f_l} w\left(p-1, f_1, \ldots, f_K\right) \quad \cdots} \quad \cdots \\
&\cdots \quad \frac{\cdots \quad + \sum_{k=1}^{K} \frac{f_k}{c-p-\sum_{l=1}^{k-1} f_l} \prod_{n=k+1}^{K} \frac{c-p-\sum_{l=1}^{n-1} f_l}{c-p-\sum_{l=1}^{n} f_l} w\left(p, f_1, \ldots, f_k - 1, \ldots, f_K\right)}{\cdots \quad + \frac{p}{c}\sum_{k=1}^{K} \prod_{n=1}^{k-1} \frac{c-p-\sum_{l=1}^{n-1} f_l}{c-p-\sum_{l=1}^{n} f_l} \frac{f_k}{c-p-\sum_{l=1}^{k} f_l} \prod_{n=k+1}^{K} \left(\frac{c-p-\sum_{l=1}^{n-1} f_l}{c-p-\sum_{l=1}^{n} f_l}\right)^2 w\left(p-1, f_1, \ldots, f_k - 1, \ldots, f_K\right)} \\
&\approx \frac{c}{p} \frac{w\left(p, f_1, \ldots, f_K\right) + \sum_{k=1}^{K} \frac{F_k}{C} w\left(p, f_1, \ldots, f_k - 1, \ldots, f_K\right)}{w\left(p-1, f_1, \ldots, f_K\right) + \sum_{k=1}^{K} \frac{F_k}{C} w\left(p-1, f_1, \ldots, f_k - 1, \ldots, f_K\right)} \\
&= \frac{c}{p} \frac{1 + \sum_{k=1}^{K} \frac{f_k}{c} \exp\left(-\beta\left(\varepsilon_{f_k} - \varepsilon_{c,f_k}\right)\right)}{\exp\left(-\beta\left(\varepsilon_p - \varepsilon_{c,p}\right)\right) + \sum_{k=1}^{K} \frac{f_k}{c} \exp\left(-\beta\left(\varepsilon_{p,f_k} - \varepsilon_{c,p} - \varepsilon_{c,f_k}\right)\right)} \\
&= \frac{c}{p} \exp\left(-\beta\left(\varepsilon_{c,p} - \varepsilon_p\right)\right) \frac{1 + \sum_{k=1}^{K} \frac{f_k}{c} \exp\left(-\beta\left(\varepsilon_{f_k} - \varepsilon_{c,f_k}\right)\right)}{1 + \sum_{k=1}^{K} \frac{f_k}{c} \exp\left(-\beta\left(\varepsilon_{p,f_k} - \varepsilon_p - \varepsilon_{c,f_k}\right)\right)}
\end{aligned}$$

Thus, we again obtain a formula for the NOT gate with imperfect competitivity and arbitrary crosstalk from other TF's:

$$\mathsf{E}\left(X \mid f_1, \ldots, f_K\right) = \frac{1}{1 + \frac{1}{d}\frac{c}{p}\exp\left(-\beta\left(\varepsilon_{c,p} - \varepsilon_p\right)\right)}$$

where the d factor is given by

$$d = \frac{1 + \sum_{k=1}^{K} \frac{f_k}{c} \exp\left(-\beta\left(\varepsilon_{p,f_k} - \varepsilon_p - \varepsilon_{c,f_k}\right)\right)}{1 + \sum_{k=1}^{K} \frac{f_k}{c} \exp\left(-\beta\left(\varepsilon_{f_k} - \varepsilon_{c,f_k}\right)\right)}$$

It is now a rather simple step to derive the final expression for the (N-input) NOR gate.

### C Genetic Gate Library Compatibility

The following figures detail the compatibility analysis of the used gate library from Cello.

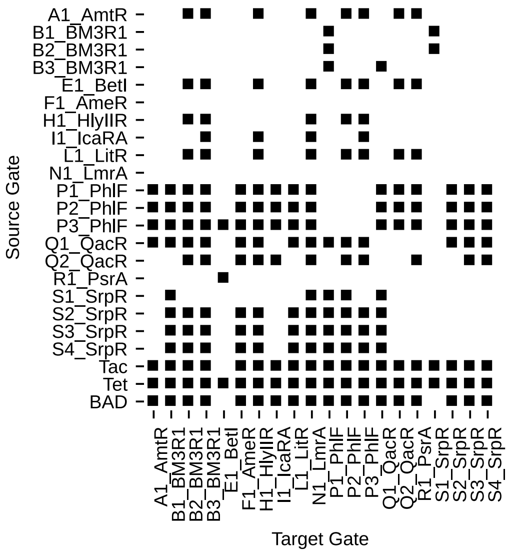


Figure 2: Pair-wise compatibility of gates according to the proposed compatibility constraint. Each dot represents a compatible pair of gates. This matrix is used for determining the compatibility of gates in the case of gates with only one input, i.e. NOT gates.

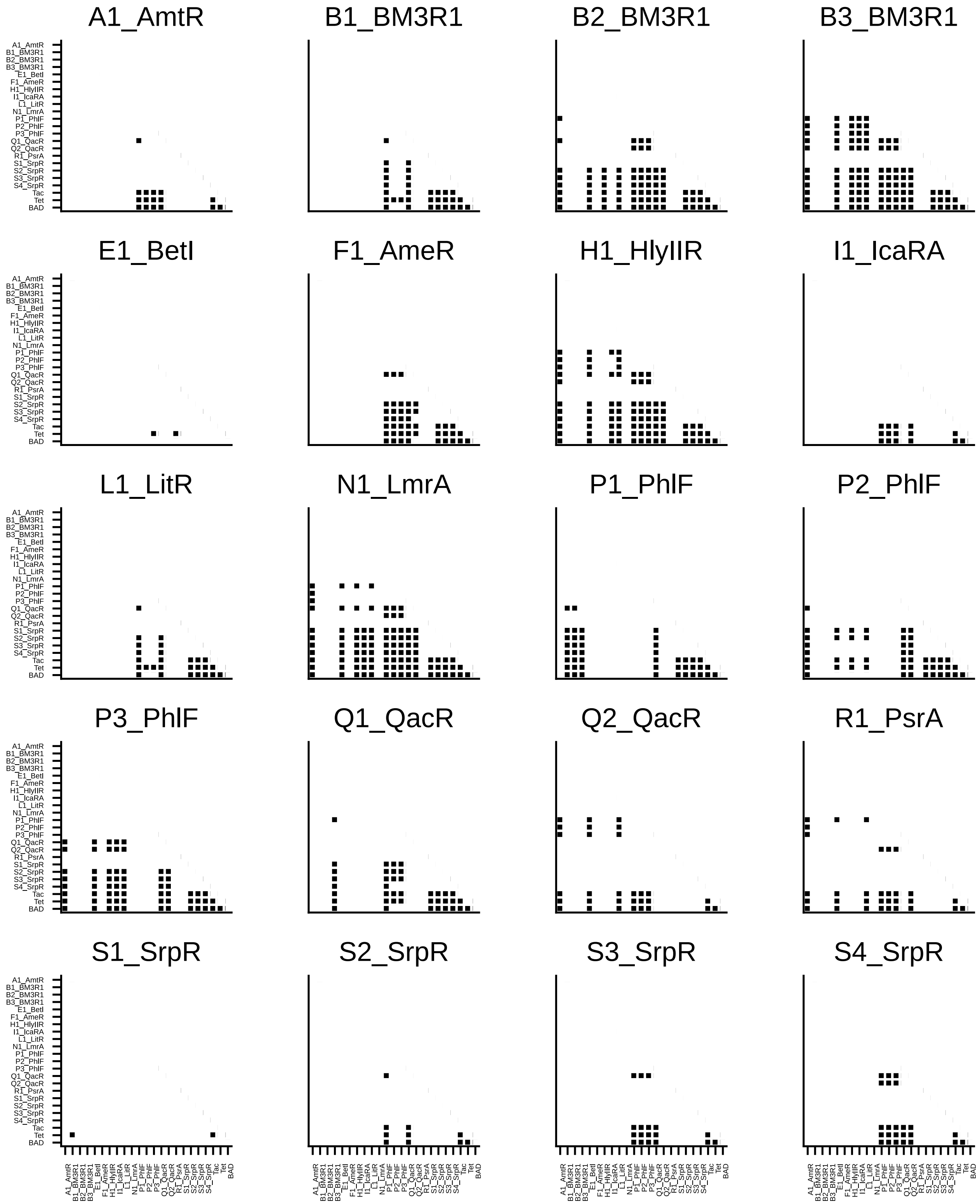


Figure 3: Detailed visualization of compatible gate triples. Each matrix shows the compatible pairs of input gates for one target gate.
